# PhosR enables processing and functional analysis of phosphoproteomic data

**DOI:** 10.1101/2020.08.31.276329

**Authors:** Hani Jieun Kim, Taiyun Kim, Nolan J Hoffman, Di Xiao, David E James, Sean J Humphrey, Pengyi Yang

**Affiliations:** School of Mathematics and Statistics, The University of Sydney, Sydney, NSW, Australia; Computational Systems Biology Group, Children’s Medical Research Institute, Faculty of Medicine and Health, The University of Sydney, Westmead, NSW, Australia; Charles Perkins Centre, The University of Sydney, Sydney, NSW, Australia; School of Environmental and Life Sciences, The University of Sydney, Sydney, NSW, Australia; Exercise and Nutrition Research Program, Mary MacKillop Institute for Health Research, Australian Catholic University, Melbourne, VIC, Australia

## Abstract

Mass spectrometry (MS)-based phosphoproteomics has revolutionised our ability to profile phosphorylation-based signalling in cells and tissues on a global scale. To infer the action of kinases and signalling pathways in phosphoproteomic experiments, we present PhosR, a set of tools and methodologies implemented in a suite of R packages facilitating comprehensive analysis of phosphoproteomic data. By applying PhosR to both published and new phosphoproteomic datasets, we demonstrate capabilities in data imputation and normalisation using a novel set of ‘stably phosphorylated sites’, and in functional analysis for inferring active kinases and signalling pathways. In particular, we introduce a ‘signalome’ construction method for identifying a collection of signalling modules to summarise and visualise the interaction of kinases and their collective actions on signal transduction. Together, our data and findings demonstrate the utility of PhosR in processing and generating novel biological knowledge from MS-based phosphoproteomic data.

## INTRODUCTION

Protein phosphorylation is an essential regulatory mechanism in cellular signal transduction. Elucidating changes in phosphorylation is crucial to understand how cells sense and respond to environmental cues and perturbations (Humphrey et al., 2015a). Advances in mass spectrometry (MS)-based technologies have enabled us to quantify changes in phosphorylation of tens of thousands of phosphorylation sites in the phosphoproteome of cells (Sharma et al., 2014). While these technological advances have enabled the generation of large-scale phosphoproteomic data (Macek et al., 2009), computational methods and pipelines for phosphoproteomic data analysis remains in relative infancy, with various challenges remaining to be addressed. Upstream steps in the analysis workflow include phosphosite filtering, handling missing values, and batch effect correction (Tyanova et al., 2016). Strategies taken in each of these data processing steps can have a considerable impact on downstream analysis of kinases and signalling pathways and enable or prohibit integrated meta-analysis of multiple phosphoproteomic datasets. Beside challenges in upstream data processing, a major obstacle in phosphoproteomics is the lack of annotated phosphosites (Needham et al. 2019). Without knowledge of cognate kinase(s) for the majority of phosphosites sites, the identification of regulated phosphosites by themselves provides an incomplete view of signalling network function. Moreover, most phosphoproteomic studies still rely on an analysis framework where phosphorylation is evaluated site-specifically although studies have revealed that many proteins are phosphorylated at multiple sites, some of which are targeted by orthogonal kinases. Adopting a phosphosite-centric analysis would therefore ignore any interactions and relationships between phosphosites from the same protein and any co-regulation of proteins at multiple sites.

Currently, only a handful of computational tools are suited to processing and downstream analysis of phosphoproteomic data. For example, whilst a large number of imputation algorithms have been developed for proteomic data (Webb-Robertson et al., 2015), significantly fewer methods are available for phosphoproteomic data (Tyanova et al., 2016). Similarly, a variety of methods developed for normalisation and batch effect correction of genomic and transcriptomic data (Johnson et al., 2007; Risso et al., 2014) have been used for phosphoproteomic data normalisation, but very few methods are specifically tailored for this task. For the downstream analysis of phosphoproteomic data, Inference of Kinase Activities from Phosphoproteomics (IKAP) (Mischnik et al., 2016), Kinase Set Enrichment Analysis (KSEA) (Casado et al., 2013), and Integrative Inferred Kinase Activity (INKA) (Beekhof et al., 2019) use kinase-substrate annotations to infer the activity of a kinases by evaluating the phosphorylation status of its substrates. However, these tools rely on a limited number of kinase-substrate relationships predicted or curated in databases such PhosphoSitePlus (Hornbeck et al., 2012) and Phospho.ELM (Dinkel et al., 2011), and may therefore restrict insight that could be obtain from unannotated sites. Although other methods including KinasePhos (Wong et al., 2007) and KinomeXplorer (Horn et al., 2014) could be used in conjuncture to address these limitations by predicting kinase-substrate relationships in a phosphoproteomic data agnostic manner, their use of motifs or protein-protein interaction network overlooks the dynamic and context-specific nature of phosphorylation. Compared to these approaches, more recent methods such as PHOTON (Rudolph et al., 2016) and CoPhosK (Ayati et al., 2019) employ a data-driven approach to infer phosphorylation-based networks. These methods enable us to begin addressing issues related to specificity of kinase-substrate relationships and context-specific regulation, but do not currently take into account how a set of phosphosites may be co-regulated within and across proteins.

Here, we developed a phosphoproteomic analysis pipeline called PhosR (**Figure 1A**) to address key issues in processing and downstream analysis of large-scale phosphoproteomic data and applied the components of PhosR to a panel of published and new skeletal muscle cell phosphoproteomic datasets. We demonstrate the impact of imputation on downstream analysis; introduce ‘stably phosphorylated sites’ (SPSs) and highlight their utility in phosphoproteomic data normalisation and integration; and introduce a kinase-substrate scoring method that leverages the dynamic profiles of canonical substrates and through which the global relationships of kinases and substrates can be annotated. We then use these annotations (1) to identify cognate proteins with phosphosites of similar regulatory profiles by interpreting phosphorylation sites in the context of their protein of origin and (2) to construct ‘signalomes’ of large-scale kinase and substrate relationships on the basis of these protein modules, and by doing so reconstruct the interactions between kinases and their collective action on signal transduction pathways. Using our approach, we demonstrate six distinct modules of cognate proteins that characterise the response of L6 myotubes to treatment with the 5’ AMP-activated protein kinase (AMPK) agonist 5-aminoimidazole-4-carboxamide-1-β-D-ribofuranoside (AICAR) and/or insulin stimulation. In particular, our approach revealed a module that is predominantly regulated by AMPK and is characterised by AMPK-activated phosphosites whose phosphorylation is attenuated with concurrent insulin stimulation. This reveals previously unappreciated interactions between the AMPK and insulin signalling pathways involved in coordinating skeletal muscle glucose transport and insulin sensitivity. Together, our data and findings demonstrate the utility of PhosR from various aspects of phosphoproteomic data processing towards generating novel biological insight from MS-based phosphoproteomic data and facilitating deeper understanding of existing and future large-scale phosphoproteomic resources.

**Figure 1.**
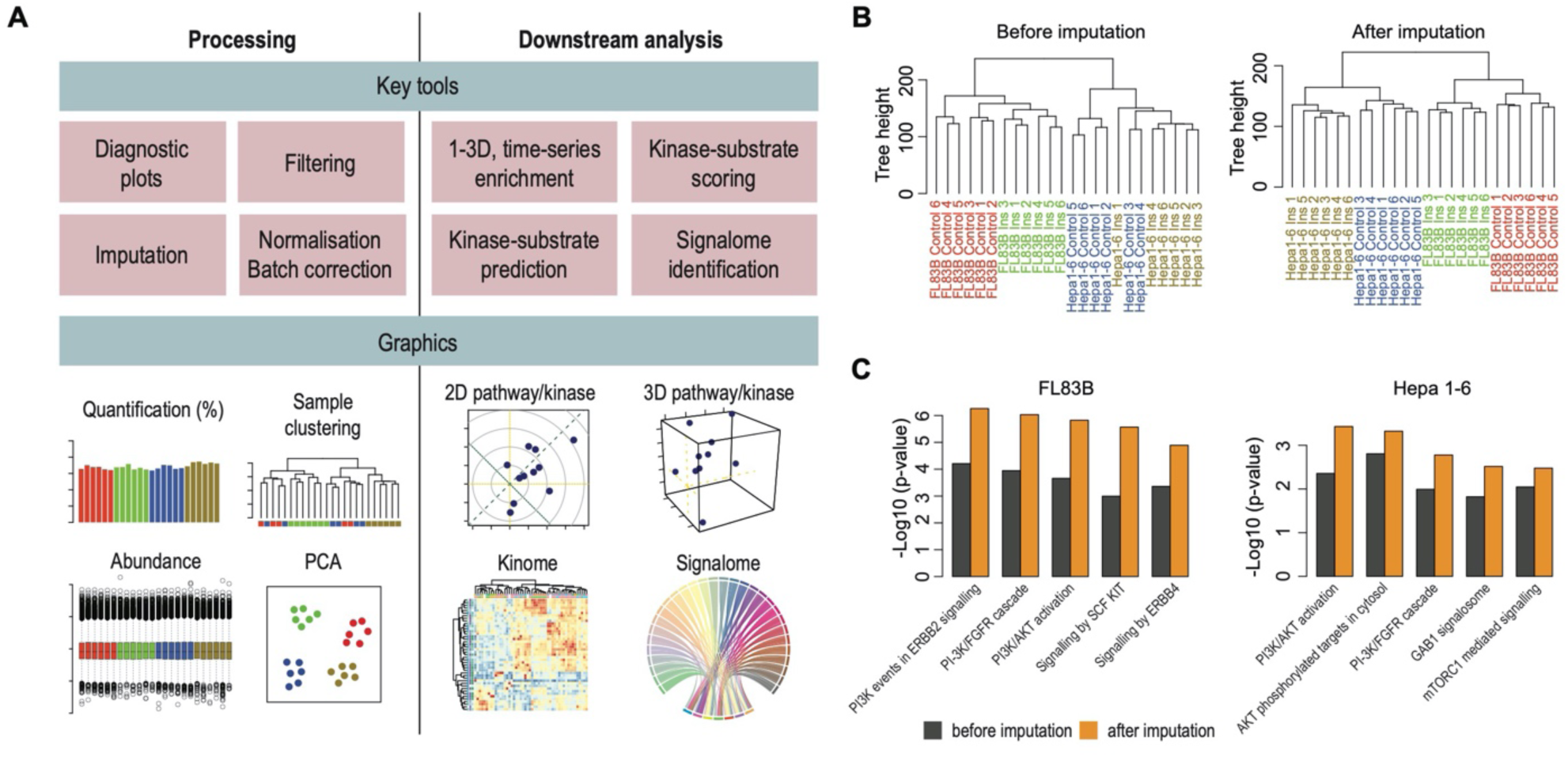
Overview of the main components of PhosR and impact on downstream phosphoproteomic analysis. (**A**) Key modules in PhosR. Modules are categorised into two broad steps of data analytics–processing and downstream analysis. PhosR implements several processing steps specifically for phosphoproteomic data, including filtering, imputation, and normalisation. Downstream analytical tools in PhosR consists of 1-, 2-, and 3-dimensional pathway analysis, kinase enrichment analysis, kinase-substrate scoring, and signalome construction. (**B**) Hierarchical clustering of biological replicates from phosphoproteomic experiments profiling FL83B and Hepa1-6 liver cells under basal or insulin-stimulated conditions. Hierarchical clustering was performed before and after imputation. (**C**) Bar plots showing enrichment of various signalling pathways known to be associated with insulin signalling before (black) and after (orange) imputation.

## RESULTS

### Pre-processing phosphoproteomic data with PhosR strengthens biological signal for downstream analysis

To first demonstrate the handling of missing data in PhosR, we used a phosphoproteome profiling dataset from FL83B and Hepa 1-6 cells stimulated with insulin (Humphrey et al., 2015b). We conducted a stepwise imputation approach, comprising site- and condition-specific imputation (*scImpute*) and paired-tail imputation (*ptImpute*) which takes into account the value of quantified phosphosites as well as the experimental design (see Methods). We demonstrate that with imputation the percentage of quantified phosphosites rises from approximately 60-75% to 80% across all samples (**Figure S1A**), and the clustering of samples showed a clear improvement relative to previous analysis when Hepa1-6 samples from different conditions were erroneously clustered together (**Figure 1B**). Next, to evaluate the impact of imputation on downstream analysis, we compared the number of differentially phosphorylated sites before and after imputation (**Figure S1B**). Interestingly, we found that while the number of differentially up- and down-regulated sites nearly doubled in FL84B cells, imputation also led to an almost three-fold decrease in the number of up-regulated sites in Hep 1-6 cells, bringing the up- and down-regulated sites to a relatively comparable range (**Figure S1B**). Using pathway enrichment analysis on the phosphoproteome summarised to protein-level with the *phosCollapse* function in PhosR (see Methods), we further show that the differentially phosphorylated sites from the imputed datasets are more enriched for the key pathways related to insulin signalling (**Figure 1C**). Together, these findings suggest PhosR imputation strengthens biological signals and facilitates downstream pathway analysis.

### Identification of a set of stably phosphorylated sites from phosphoproteomic data

Several commonly used data normalisation approaches such as the “removal of unwanted variation” (RUV) (Gagnon-Bartsch and Speed, 2012) require a set of internal standards that are known to be unchanged biologically in the samples measured. This is a challenge for phosphoproteomics since phosphorylation is highly dynamic, with diverse regulation across different cell types and experimental conditions. To explore whether we could identify a set of phosphorylation sites that might meet the criteria of being “stably phosphorylated” across multiple phosphoproteomics datasets we used four high-quality datasets generated from different cell types, and experimental conditions. These include the phosphoproteome data from the time-course of mouse embryonic stem cell differentiation (Yang et al., 2019) (‘ESC differentiation’), phosphoproteomic data of control and FGF21 treated mouse 3T3-L1 adipocytes (Minard et al., 2016) (‘Adipocyte FGF21’), phosphoproteomic data of FL83B and Hep 1-6 cells (‘FL83B & Hep 1-6 insulin’; processed as described in the previous section), and phosphoproteomes of control and insulin stimulated, and kinase inhibitor treated mouse 3T3-L1 adipocytes (Humphrey et al., 2013) (‘Adipocyte insulin, LY, & MK’). We performed a four-way overlap of the four datasets and found 1,207 phosphosites common to all four datasets (**Figure S2A**). To identify stably phosphorylated sites, we ranked the overlapping phosphosites on the basis of their absolute log2 fold change (**Figure S2B**) and generated a consensus ranking using a statistical framework (see Methods) where phosphosites with consistently small fold changes are highly ranked and those with large fold changes are lowly ranked (**Figure S2C-D**). The top 100 phosphosites from the consensus list are referred hereafter as stably phosphorylated sites (SPSs) and are included in the PhosR package as a resource for normalisation of future phosphoproteomic data.

### Normalisation using stably phosphorylated sites removes unwanted variation in phosphoproteomic data

To evaluate the utility of SPSs in phosphoproteomic data normalisation, we applied RUV-III (Molania et al., 2019) with SPSs (denoted as ‘*RUVphospho*’) to normalise the phosphoproteomic data of L6 myotubes treated with AICAR, an analog of adenosine monophosphate (AMP) that stimulates AMPK activity, and insulin either individually or in combination (**Figure 2A**; see Methods). Before normalisation, hierarchical clustering and principal component analysis (PCA) of the myotube phosphoproteomic data revealed a batch effect that is driven by experiment runs for samples treated with insulin (**Figure 2B, C**; left panels), whereas normalisation with *RUVphospho* effectively corrects this (**Figure 2B-C**; right panels).

**Figure 2.**
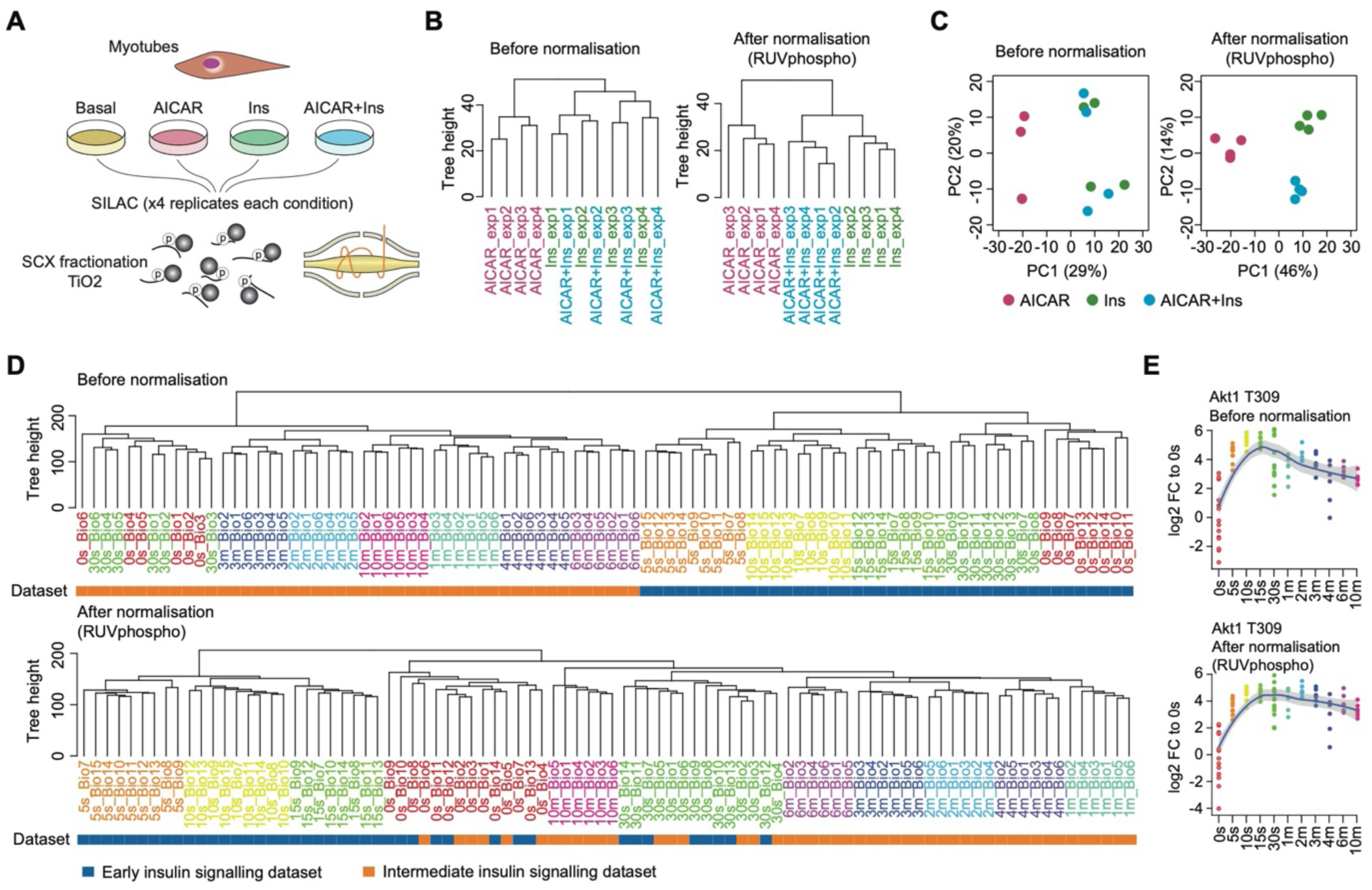
Normalisation and batch correction using RUV and SPSs in PhosR. (**A**) Schematic illustrating the experimental setup of the phosphoproteomic profiling experiment in L6 myotubes where phosphoproteomic analysis of cells were performed in the basal condition or following treatment with the AMPK agonist AICAR, insulin (Ins), or in combination (AICAR+Ins). (**B**) Dendrogram of all biological replicates before and after *RUVphospho* normalisation of the myotube phosphoproteomic data. Samples are coloured by experimental condition. (**C**) PCA plot of the myotube phosphoproteomes before and after *RUVphospho* normalisation. Each point represents a sample and is coloured by experimental condition. (**D**) Hierarchical clustering of phosphoproteomic datasets from early and intermediate in situ insulin stimulation of mouse liver before and after *RUVphospho* normalisation. Samples are coloured by timepoint and dataset. (**E**) Scatter plot comparing the log2 fold-change in phosphorylation of AKT1 T309 upon insulin stimulation before and after implementation of PhosR normalisation.

Integrating multiple phosphoproteomics datasets from independent studies is typically challenging, since signal derived from technical sources such as HPLC and mass spectrometer performance characteristics often dominate biological signals. To illustrate the ability of *RUVphospho* in enabling the integration of phosphoproteomics data from independent studies, we used two time-course datasets from early and intermediate insulin signalling from in mouse liver (Humphrey et al., 2015b). This study was not included amongst the 4 used to select the stably phosphorylated sites. It contains two overlapping time points in each time-series (0s and 30s), thus providing the opportunity to integrate the time-series into a single comprehensive dataset. Prior to normalization, hierarchical clustering of the combined datasets reveals separation of the independent time-series. Applying *RUVphospho* effectively integrates the two datasets, as demonstrated by the clustering of 0 and 30s timepoint samples from the two datasets (**Figure 2D**). Closer inspection of the temporal phosphorylation change of phosphosite AKT1 T309, one of the most important markers of AKT activity in response to insulin stimulation (Humphrey and James, 2012), reveals a smoother temporal profile following normalisation with *RUVphospho*. Collectively, these results demonstrate the normalisation procedure in PhosR facilitates effective batch correction and integration of phosphoproteomic data.

### Dual-centric analyses to detect regulated pathways and kinases in phosphoproteomic data

Most phosphoproteomic studies have adopted a phosphosite-level analysis of the data. To enable phosphoproteomic data analysis on the protein level, PhosR implements both site- and protein-centric analyses for detecting changes in kinase activities and signalling pathways through traditional enrichment analyses (over-representation or rank-based gene set test, together referred to as ‘1-dimensional enrichment analysis’) as well as 2- and 3-dimensional analyses (**Figure 1A**). To test which signalling pathways are activated upon insulin stimulation in myotubes, we performed protein-centric enrichment analyses on the normalised myotube phosphoproteomic dataset using both over-representation and rank-based gene set tests (**Figure S3A**). We found several expected pathways including those associated with mTORC1, AKT, and ERK signalling. While these highly enriched pathways were largely in agreement between the two types of enrichment analyses, the rank-based gene set test had much greater statistical power in detecting these pathways (**Figure S3B**).

The 2- and 3-dimensional analyses implemented in PhosR use direction-based statistics (Yang et al., 2014, 2016) which enables the investigation of kinases regulated by different combinations of treatments. Applying this to the myotube phosphoproteome datasets, we found that, as expected, the activity of AMPK, marked by PRKAA1 (the catalytic alpha-1 subunit of AMPK), is up-regulated following both AICAR and AICAR+Ins treatments but remains unchanged by insulin treatment alone (**Figure S3C**; top two panels). Strikingly, we found that the AICAR-induced up-regulation of AMPK catalytic activity is attenuated by the addition of insulin as is observed from the kinase activity plot of AICAR versus AICAR+Ins (**Figure S3C**; bottom left panel). These pairwise comparisons can be summarised using the 3D analysis where the three comparisons are integrated into a single statistical analysis to highlight the combinatorial effect of different treatments on PRKAA1 activity (**Figure S3D**).

### Global kinase-substrate relationship scoring of phosphosites using PhosR

A key challenge in analysing phosphoproteomics data is in identifying kinases responsible for the phosphorylation of specific sites. While various computational tools can be applied to annotate potential kinases of particular phosphosites identified in the phosphoproteomic data on the basis of their amino acid sequences or structural information (Trost and Kusalik, 2011), most methods do not directly consider cell type and/or treatment/condition specificity of phosphorylation. To this end, PhosR implements a multi-step kinase-substrate scoring method where first the likelihood of a kinase to regulate a phosphosite is scored by combining both kinase recognition motifs and the dynamic phosphorylation profiles of sites. The combined scores across all kinases are then integrated using an adaptive sampling-based positive-unlabelled learning method (Yang et al., 2018) to prioritise the kinase most likely to regulate a phosphosite (see Methods). The application of the proposed scoring method to the myotube phosphoproteome uncovers potential kinase-substrate pairs (**Figure 3A**; row dendrogram) and global relationships between kinases (**Figure 3A**; column dendrogram). A KEGG pathway overrepresentation analysis of the kinase-substrate pairs highlights overrepresented pathways known to be associated with each kinase (**Figure S4A**). The kinase dendrogram reveals three major kinase groups governing the myotube phosphoproteome (AGC kinases [e.g. RPS6K and AKT isoforms], CMGC kinases [e.g. MAPKs], and CAMK kinases [e.g. AMPK catalytic subunits PRKAA1 and PRKAA2]). In particular, our kinase-substrate scoring method confirms several well-established substrates of AMPK such as ACACA S79, AKAP1 S103, SMCR8 S488 (Hoffman et al., 2015), and MTFR1L S100 (Schaffer et al., 2015) while also finding new candidate AMPK substrates such as XIRP1 S532 and MAP4K4 S791 (**Figure 3A, B and Figure S4B**). Importantly, except for ACACA S79, the above-mentioned phosphosites are not annotated as substrates of AMPK in the PhosphoSitePlus database. In agreement with our 2D kinase enrichment analysis (**Figure S3C**; bottom left panel), we demonstrate that the phosphorylation profiles of several of these AMPK substrates show a strong up-regulation of phosphorylation upon AICAR stimulation that is attenuated when myotubes are co-stimulated with insulin (**Figure 3C**).

**Figure 3.**
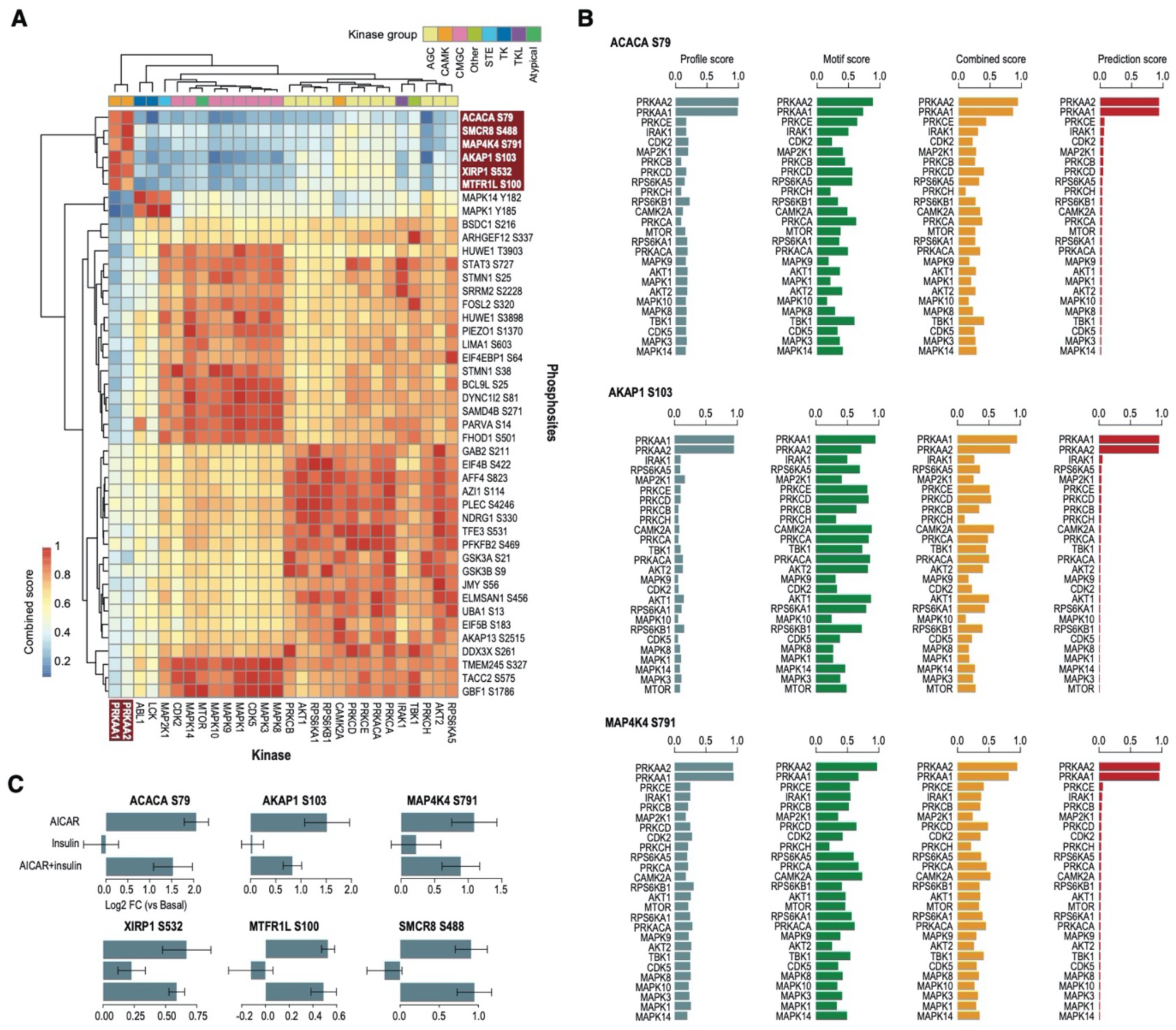
Global kinase-substrate relationship scoring of the myotube phosphoproteome. (**A**) A clustered heatmap of the combined score from kinase-substrate scoring function for the top three phosphosites of all kinases evaluated in this study. A higher combined score denotes a better fit to a kinase motif and kinase-substrate phosphorylation profile of a phosphosite. Kinases are annotated to which kinase group they belong. (**B**) Bar plots showing profile, motif, and combined scores, and positive-unlabelled ensemble learning prediction score of the top-ranked AMPK substrates. (**C**) Bar plots showing the log2 fold change (Log2 FC) in phosphorylation level of the top-ranked AMPK substrates after treatment with AICAR, insulin, and combined treatment.

### Construction of signalomes from discrete modules of co-regulated proteins

Proteins are frequently phosphorylated at multiple sites and often by orthogonal kinases. Site- and protein-centric analyses of phosphoproteomics data lie at opposite ends of the spectrum, with the former treating phosphosites on the same protein independently and ignoring the host protein information, and the latter focussing on a specific site, losing information from other sites on the same protein. Because of the lack of appropriate methods, the question of whether proteins are co-regulated across multiple phosphosites remains poorly investigated. Leveraging our global kinase-substrate scoring of phosphosites, we set out to generate signalomes wherein dynamic changes in phosphorylation within and across proteins are conjointly analysed.

We developed an approach to generate signalomes comprising discrete protein modules with phosphosites sharing similar dynamic phosphorylation profiles and kinase regulation (**Figure 4A**; see Methods). Using this approach, we show that the myotube phosphoproteome stimulated by AMPK activation and/or insulin stimulation contains six discrete protein modules. The resulting map of signalomes demonstrates that the modules are regulated by different kinases and at varying proportions (**Figure 4B**). Notably, the signalome map highlights a module (blue, module 3) entirely regulated by AMPK catalytic activity (PRKAA1 and PRKAA2) and others (orange, module 1; and green, module 4) that are co-regulated by AMPK with other kinases suggesting potential signalling crosstalk (**Figure 4B**). We then zoomed in to the extended AMPK signalome (see Methods) from the signalome network (**Figure 4C**) and found distinct phosphorylation profiles between the three protein modules (**Figure 4D and S5B)**. Consistent with previous reports (Kjøbsted et al., 2015), the phosphorylation of sites in modules 1 and 4 show synergistic effects upon AICAR and insulin stimulation. Yet in agreement with our kinase activity analysis, module 3 – predominately regulated by AMPK alone – displays activity that is enhanced by AICAR and attenuated by insulin (**Figure 4D**).

**Figure 4.**
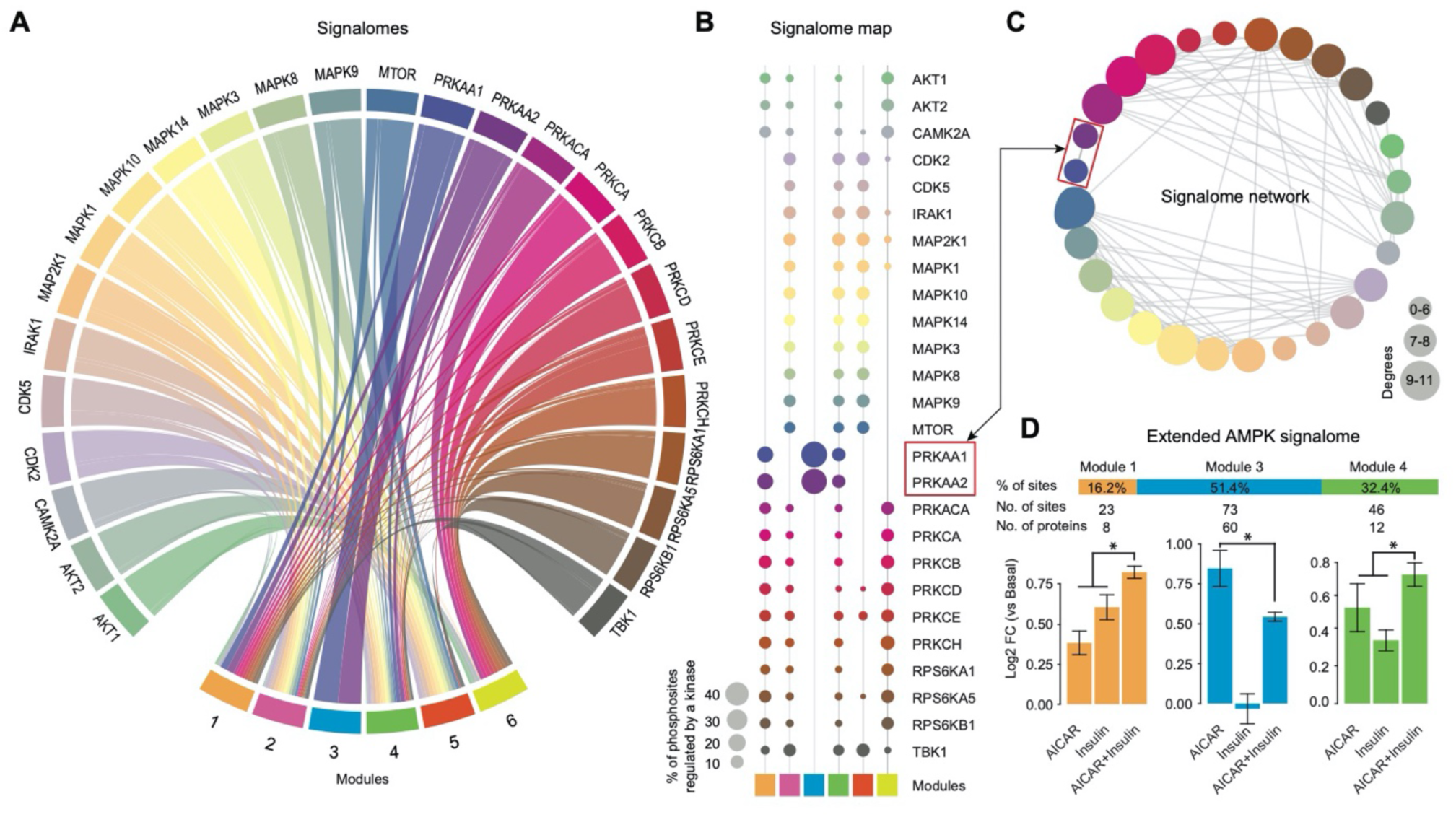
Construction of signalomes in the myotube phosphoproteome. (**A**) A chord diagram of the signalomes identified from the myotube phosphoproteome. The branching nodes consist of 26 kinases and the stem nodes consist of 6 protein modules each with a distinct phosphorylation and regulatory profile. Edges between nodes connect kinases to the protein modules they regulate. (**B**) The signalome map demonstrating the proportion of phosphosites regulated by kinases in each protein module. The size of the balloon denotes the percentage of phosphosites the kinase regulates within a module. (**C**) An interaction network of kinases. The higher the number of interactions with other kinases (degrees), the larger the circle. (**D**) Summary of the three protein modules present in the extended AMPK signalomes. The proportion of phosphosites, the total number of phosphosites, and the total number of proteins comprising the modules in the signalome is summarised. Bar plots of log2 foldchange in phosphorylation of the phosphosites in each module is summarised for each condition against basal. The error bars denote standard deviation and *indicates a p<0.05 using Wilcox rank sum test.

## DISCUSSION

Here we present PhosR, a complete set of methods and tools for phosphoproteomic data processing and downstream analysis. Using PhosR, we have at once addressed many current challenges facing phosphoproteomic data analysis. We have addressed issues of data imputation, normalisation, and integration through *RUVphospho*, supported by defining a set of stably phosphorylated sites (SPSs), which we include in the PhosR package as a resource to the community. Processing of phosphoproteomics data with PhosR facilitated the extraction of differentially phosphorylated proteins with greater biological relevance, demonstrated by strengthened signal of known pathways. *RUVphospho* normalisation enabled datasets from independent studies to be integrated and elimination of batch effects without affecting biological signal.

A major challenge in phosphoproteomics studies is the lack of annotated phosphosites, and with less than 5% of the phosphoproteome with a known functional link to a kinase, our ability to glean functional interpretation from phosphoproteomic data is severely limited (Needham et al., 2019). Biochemically assigning phosphosites to their cognate kinase is an experimentally labour-intensive process and may be affected by the experimental system employed. Moreover, because of the great complexity within phosphoproteomes, many kinase-substrate relationships are likely to be context and cell-type specific, further complicating efforts to elucidate them. Our global kinase-substrate scoring method enables the inference of kinase activities specific to experimental conditions and cellular systems and the construction of signalomes wherein both dynamic and differential phosphorylation changes in phosphosites within and across proteins are taken into account. Using this approach, we could identify proteins co-regulated at three levels: 1) across experimental conditions; 2) between multiple phosphosites; and 3) by similar kinase regulation. In doing so, we can begin investigating these layers of complexities in signal transduction networks.

Previous research has demonstrated that skeletal muscle AMPK activation, for example following AICAR treatment or exercise, influences muscle glucose transport and insulin sensitivity and is therefore an important regulator of glucose homeostasis and metabolic health. In particular, prior stimulation of muscle skeletal muscle with AICAR to stimulate AMPK activity has been shown to lead to enhance the sensitivity with which insulin stimulates glucose uptake (Kjøbsted et al., 2015). Our approach to generate modules of co-regulated proteins enabled the discovery of three sets of proteins with phosphosites that are regulated by AMPK in the myotube phosphoproteome stimulated with either AICAR, insulin, or in combination (**Figure 4A, B**). Consistent with previous knowledge, we found two modules (modules 1 and 4) that exhibited enhanced phosphorylation upon insulin treatment if they were first stimulated by AICAR, demonstrating a synergistic effect between insulin and AMPK signalling pathways in these modules (**Figure 4D**). Indeed, in module 1 we observed TBC1D1, which has been implicated in the AMPK-dependent increase of muscle glucose uptake and insulin sensitivity (Dokas et al., 2013; Kjøbsted et al., 2015; Taylor et al., 2008). Intriguingly, our approach also revealed a module (module 3) entirely regulated by AMPK and, unlike the other two modules, the phosphosites found in this protein module demonstrated strong activation by AICAR treatment and no sensitivity to insulin stimulation alone. Strikingly, the AICAR induced activation of phosphorylation on these sites was attenuated by the addition of insulin, suggesting a negative regulatory effect of insulin on the phosphorylation of AMPK substrates (**Figure 4D**). Because key differences between module 3 and modules 1 and 4 are differential kinase regulation of phosphosites and the presence of insulin-sensitive sites, we postulate that the interplay of AMPK with other kinases such as MAPKs and S6K may occur to stimulate diverse actions on different signalling pathways. In conclusion, our signalome construction method is applicable to diverse datasets that profile dynamic changes in the phosphoproteomes, enables inference of kinase activities through new visualisation of kinase interactions and their collective action on signal transduction pathways, and supporting the interpretation of phosphoproteomic data at a level beyond the analysis of phosphosites in isolation.

## METHODS

### Experimental Data

This study utilised a collection of published phosphoproteomic datasets and profiled the phosphoproteome of myotubes in response to AICAR and insulin stimulation (new data). The published datasets used in this study include (i) the ‘ESC differentiation’ dataset where a cocktail of treatments were applied to differentiate mouse embryonic stem cells (ESCs) to epiblast-like cells (Yang et al., 2019) (PRIDE: PXD010621). The phosphoproteome of this ESC differentiation process was profiled using label-free quantification at 12 selected time points; (ii) the ‘Adipocyte FGF21’ dataset (Minard et al., 2016) (PXD003631) where the phosphoproteome of 3T3-L1 adipocytes treated with either insulin or FGF21 were profiled using SILAC quantification; (iii) the ‘Adipocyte insulin, LY, and MK’ dataset (Humphrey et al., 2013) where the phosphoproteome of 3T3-L1 adipocytes treated with either insulin in a time-course or LY and MK prior to insulin were profiled using SILAC quantification; (iiii) the ‘FL83B and Hepa 1-6 Insulin’ dataset (Humphrey et al., 2015b) (PXD001792) where the phosphoproteome of FL83B and Hepa 1-6 cells in basal and treated with insulin were profiled using label-free quantification; and (iv) the ‘Mouse liver insulin’ datasets (Humphrey et al., 2015b) (PXD001792) where the phosphoproteomes of mouse livers treated with insulin were profiled in an early time-course and an intermediate time-course using label-free quantification.

For profiling the phosphoproteome response of rat L6 myotubes to AMPK activation and insulin stimulation, stable isotope labelling by amino acids (SILAC)-based quantification (Ong et al., 2002) was performed in biological quadruplicates on cells in basal condition or treated with AICAR (2 mM, 30 min) and insulin (100 nM, 20 min) individually or in combination. Following cell harvesting and mixing of equal protein content from light and heavy SILAC cell populations, proteins were trypsinised and fractionated using strong cation exchange chromatography and phosphopetides were enriched and analysed by LC-MS/MS as described previously (Hoffman et al., 2015). Raw MS data were processed using MaxQuant (Cox and Mann, 2008) (version 1.5) by searching with the following variable modifications: methionine oxidation; and serine, threonine and tyrosine phosphorylation. The precursor-ion mass tolerance was set to 20 ppm and 7 pm for first and second searches respectively and product-ion mass tolerance set to 0.02 Da. An FDR cutoff of 0.01 was used at the peptide level for selecting high confidence peptide identifications.

### Phosphosite Filtering and Imputation

MS-based phosphoproteomic data commonly contain a large amount of missing values due to biological and technical reasons. PhosR implements a collection of data filtering and imputation methods for dealing with missing values in a phosphoproteomic dataset. For filtering, PhosR allows users to specify an overall quantification rate (i.e. the percentage of quantification) of a phosphosite across all biological replicates of all conditions (or time points in a time-course experiment) from which phosphosites with lower quantification rate would be removed from further analysis. While this overall quantification rate filtering is straightforward to implement, more flexible filtering procedures are needed in many scenarios. One common scenario is that a phosphosite is only phosphorylated in a specific condition (or treatment) but not in other conditions. Let us denote the quantification rate of a phosphosite in biological replicates of a condition as *q*^*t*^(*t =* 1 … *T*) where *T* is the number of conditions (or time points in the case of time-course data). PhosR implements the function *selectGrps* which allows phosphosites with a *q*^*t*^ value equal to or greater than a predefined threshold in one or more conditions to be retained. A schematic example is shown in **Figure S6A**. In addition, for time-course data, PhosR implements the function *selectTimes* which allows phosphosites with a *q*^*t*^ value equal or greater than a predefined threshold in two or more consecutive time points to be retained.

For data imputation, PhosR implements multiple methods to take advantage of data structure and experimental design. These include site- and condition-specific imputation (*scImpute*) where, in a condition, the missing values of a phosphosite with a *q*^*t*^ value equal or greater than a predefined threshold will be imputed by sampling from the empirical normal distribution constructed from the quantified values of that phosphosite in that condition (**Figure S6B**); tail-based imputation (*tImpute*), similar to those described in (Tyanova et al., 2016), where the missing values were imputed from the tail of the empirical normal distribution with a default setting of *𝒩* (*μ* − *σ* × 1.6, *σ* × 0.6) constructed from the quantified values across all sites in a sample (**Figure S6C**); and paired tail-based imputation (*ptImpute*) where for a phosphosite that have missing values in all replicates in a condition (e.g. ‘basal’) and a *q*^*t*^ value equal or greater than a predefined threshold in another condition (e.g. ‘stimulation’), the tail-based imputation is applied to impute for the missing values in the first condition (**Figure S6D**).

### Identification of Stably Phosphorylated Sites and Data Normalisation

To identify a set of stably phosphorylated sites (SPSs) for subsequent data normalisation and batch effect correction, we utilised four independent datasets including ‘ESC differentiation’, ‘Adipocyte FGF21’, ‘Adipocyte insulin, LY, and MK’ and ‘Hepa 1-6 and FL83B insulin’ (see **Experimental Data**). We selected phosphosites that were identified in all four datasets and then applied multiple steps to rank the selected phosphosites. For each dataset, let us denote the log2 quantification of a selected phosphosite compared to a control condition as 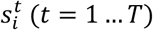 where *T* is the number of conditions or time points in that dataset. We first calculated the rank of each phosphosite by max(abs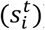) in each dataset. This captures the maximum magnitude of changes, either up- or down-regulation, of each phosphosite in each of the four datasets. We next converted the ranks of phosphosites in each dataset into z-scores from which we calculated the *p*-values *p*_*i*_ (*i =* 1, 2, 3, 4) for each phosphosite in each of the four datasets. The *p*-values of each phosphosite were then integrated into a single combined *p*-value using Fisher’s methods:

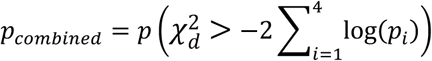

The *p*_*combained*_ was used to generate the final consensus ranking of phosphosites identified in all four dataset and the top-100 sites that show the overall minimum phosphorylation level changes were selected as SPSs.

To perform data normalisation and batch effect correction, we implemented a wrapper function *RUVphospho* which makes use of SPSs identified above as ‘negative controls’ in the RUV method using the version RUV-III (Molania et al., 2019). When the input data contains missing values, tail-based imputation will be applied to impute for the missing values since RUV-III requires a complete data matrix (**Figure S6E**). The imputed values are removed by default after normalisation but can be retained for downstream analysis.

### Protein- and Phosphosite-centric Enrichment Analyses

To enable enrichment analyses on both gene and phosphosite levels, PhosR implements a simple method called *phosCollapse* which reduces phosphosite level of information to the protein level by selecting the sites with either the maximum (by default) or minimum 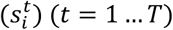 values as the representative of phosphorylation changes of their respective proteins. Phosphosite-centric analyses are performed using kinase-substrate annotation information from PhosphoSitePlus and protein-centric analyses are performed using Reactome and KEGG databases while other pathway annotation databases such as Gene Ontology can also be used as well. For testing enrichment, PhosR implements two typical methods including over-representation test (using Fisher’s Exact test) and rank-based gene set test (using Wilcoxon rank-sum test), and together refer to as 1-dimensional enrichment analyses. PhosR also provide a single interface to unify several methods developed previously for analysing multiple experimental conditions simultaneously (refer to as 2- and 3-dimensional enrichment analyses (Yang et al., 2014, 2016).

### Kinase-substrate Prioritisation of Phosphosites

To identify potential kinases that could be responsible for the phosphorylation change of a phosphorylation site, we implement a multi-step framework that contains two major components including (i) a *kinaseSubstrateScore* function which scores a given phosphosite using kinase recognition motif and phosphoproteomic dynamics, and (ii) a *kinaseSubstratePred* function which synthesise the scores generated from (i) for predicting kinase-substrate relationships using an adaptive sampling-based positive-unlabelled learning method (Yang et al., 2018). The kinase-substrate scoring function combines both kinase recognition motif (i.e. motif matching score) and experimental perturbation (i.e. profile matching score) for prioritising kinases that may be regulating the phosphorylation level of each site quantified in the dataset. To calculate the motif matching score for each kinase, all kinases and their substrate peptide sequences from PhosphoSitePlus database were used to compile position-specific scoring matrices (PSSMs) as follows:

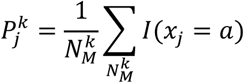

where *k* (*k =* 1 *… K*_*M*_) is the index of kinases, 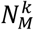 is the number of substrate sequences included for calculating the PSSM for the *k*th kinase, *j* is the index to a position in sequence *x* (with a window size of 13 surrounding the sites of phosphorylation), and *a* is the set of characters corresponding to the 22 amino acids. Then, a motif matching score is calculated for each of all phosphorylation sites *s*_*i*_ by scoring their surrounding amino acid sequences *x*_*i*_ against each of all PSSMs for quantifying the phosphorylation preference of kinases to each phosphosite:

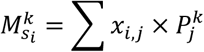

For calculating profile matching score, the phospho-quantification of each site in the phosphoproteomic data is first z-score transformed. Then, for each of all kinases, PhosR searches in the phosphoproteomic data for any known substrates of each of all kinases. For each kinase that have one or more known substrates quantified in the phosphoproteomic data 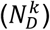, the z-score transformed dynamic phosphorylation profiles of its known substrates are median averaged (denoted as *d*_*k*_(*k =* 1 … *K*_*D*_), where *Kd* is the total number of kinases that have a quantified substrate profile). Next, the profile matching scores of each phosphosite quantified in the dataset are calculated by using Pearson’s correlation with respect to the averaged profiles of known substrates of each of all kinases:

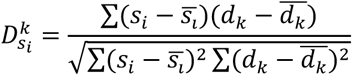

The final combined score of a phosphosite *s*_*i*_ with respect to a kinase *k* is the weighted average of the motif matching score and profile matching score by taking into the number of sequences and substrates used for calculating the motif and profile of the kinase. Specifically, the weights for the two parts of a kinase are calculated as 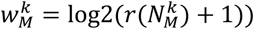 and 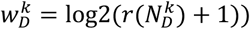 and the combined score is calculated as:

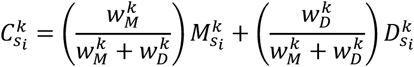

While the combined score calculated above takes into account both motif and phosphorylation profile of a phosphosite in prioritising kinases that may responsible for their phosphorylation changes, these scores for each kinase are calculated independently from each other. To maximise the information in determining kinase-substrate relationships, PhosR implements a machine learning method where the combined scores across all kinases are used as learning features to predict for kinases 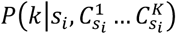 for a given phosphosite *s*_*i*_. One of the key issues in training machine learning models for predicting kinase substrates is the need to curate a set of training examples for each kinase. This is difficult for most kinases because the numbers of known substrates are prohibitively small for training predictive models. To this end, we implemented in PhosR the AdaSampling-based positive-unlabelled ensemble of support vector machines (SVMs) as described previously (Yang et al., 2018). For each kinase the top 30 highly ranked phosphosites (based on the combined scores) are used initially as positive examples for training SVMs for predicting substrates of that kinase and the AdaSampling procedure is used to subsequently update the training examples based on the model confidence on each phosphosite.

### Signalome Construction

To construct signalomes wherein kinase regulation of protein modules can be identified, we developed an approach where we make direct use of the kinase-substrate prioritisation scores from the functions *kinaseSubstrateScore* and *kinaseSubstratePred*.

A similarity matrix of phosphosites is generated from the combined score from the *kinaseSubstrateScore* function by using Pearson’s correlation as the similarity metric. The resulting matrix provides a correlation of the kinase-substrate scoring of phosphosites against all other phosphosites. The similarity matrix is then used to hierarchically cluster the phosphosites into groups with distinct profiles. Because the kinase-substrate scoring is a combined score of both kinase recognition motif (i.e. motif matching score) and experimental perturbation (i.e. profile matching score) for a phosphosite against all kinases, the phosphosites are partitioned into clusters on the basis of all these components, whilst taking into account the global relationships between kinases. The total number of phosphosite clusters is determined as the number of clusters wherein the mean correlation is equal to or above 0.5 for all clusters. When there are multiple scenarios where all clusters have an average correlation equal to or above 0.5, the set of clusters with the highest average correlation is chosen.

Given that many proteins are found to have multiple differentially regulated phosphosites, many of which were predicted to be regulated by different kinases, we devised a method to evaluate phosphoproteomic data whereby the regulation of multiple phosphosites can be analysed at the protein-level (therefore allowing both a protein- *and* site-centric analysis). To this end, we constructed a phosphosite co-assignment matrix based on the phosphosite clusters and the proteins they reside on. The co-assignment matrix essentially provides a way to assign phosphosites of each protein across the clusters, generating a profile of assignment. As proteins will show different profiles in terms of their overall phosphosite membership across the clusters, we are able to create multiple combinations of protein assignment. The assignment is a binary score, meaning that the frequency with which a protein is assigned is not considered, ensuring that the co-assignment matrix does not bias towards proteins with many phosphosites. The final co-assignments, herein referred to as “protein modules”, consist of exclusive sets of proteins with similar phosphorylation profiles and kinase regulation.

The *Signalomes* function generates a visualisation of the signalomes present in the phosphoproteomic data. For the visualisation of signalomes, it does so by using the protein modules identified from above and the kinase-substrate predictions from the *kinaseSubstratePred* function. A cut-off of 0.5 is used as default (*signalomeCutoff* = 0.5) to capture kinase-substrate relationships (**Figure S4b**). Then an adjacency matrix depicting the regulation of proteins by kinases is used to generate a chord diagram from the *circlize* package. This method of visualisation provides a summary of the kinase regulation of each protein module. The *Signalomes* function also outputs signalomes associated to any kinase of interest (referred to as extended signalome of a kinase). To facilitate assessment of proteins and phosphosites that are under similar regulation, the extended signalome of a kinase combines cognate signalomes from other kinases that share a high degree of similarity in substrate regulation.

### Differentially Phosphorylated Phosphosites

Differentially phosphorylated sites were identified using the two-sided moderated *t*-test implemented in the *limma* R package (Ritchie et al., 2015). Analyses were done on log (base 2)-transformed data and *p*-values were adjusted for multiple testing using Benjamini-Hochberge FDR correction at *α=* 0.05.

### KEGG Pathway Over-representation Analysis

Pathway over-representation analysis was performed on protein sets identified from our kinase-substrate scoring analysis (kinase-substrate pairs with score > 0.5 were selected) using the over-representation analysis implemented in the *clusterProfiler* R package (Yu et al., 2012). The KEGG pathway database was used and *p*-values were adjusted for multiple testing using Benjamini-Hochberge FDR correction at *α=* 0.05.

### Software

The PhosR package written in R programming language is available at the Github repository (https://github.com/PYangLab/PhosR). All source code is published under the open-source license of GPL-3 adhering to R open-source requirement and is freely accessible to the scientific community.

## AUTHOR CONTRIBUTIONS

P.Y. designed the computational methods with input from H.J.K. and S.J.H. H.J.K., T.K. and P.Y. performed the computational analysis. T.K., H.J.K. and P.Y. implemented the R package. D.X. assisted the computational analysis and package implementation. N.J.H and S.J.H. performed myotube sample preparation and MS analysis under the supervision of D.E.J. All authors wrote and edited the manuscript.

## SUPPLEMENTARY MATERIAL

**Figure S1.**
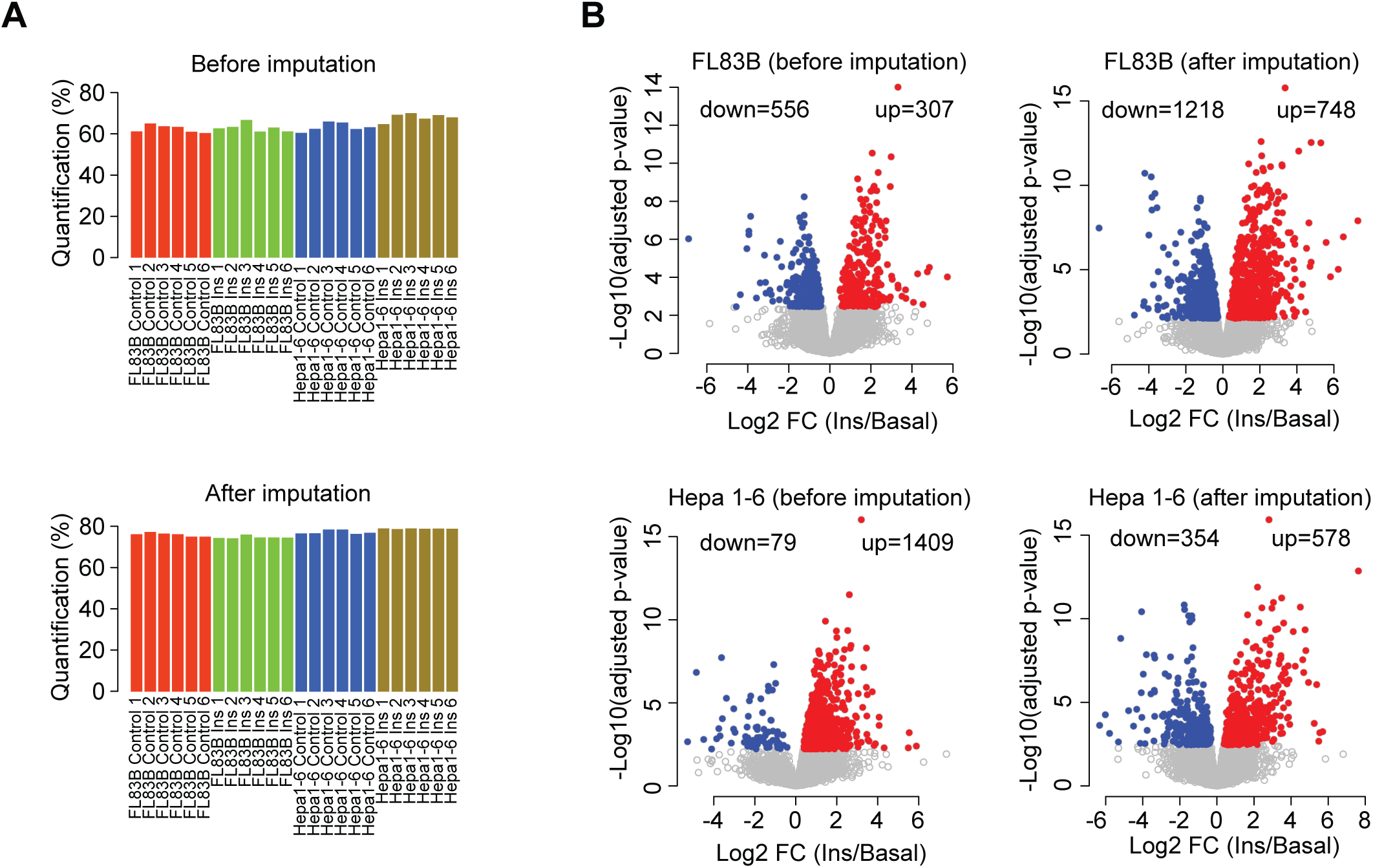
Improvement in detection of differentially expressed genes after imputation of phosphoproteomics data. (**A**) Bar plots showing the percentage of quantified sites in each sample before and after imputation. In each group, there are six biological replicates. For imputation, we used *scImpute* for phosphosites with >50% quantification of replicates in each condition and cell type and *ptImpute* for phosphosites quantified in >50% replicates in insulin treated samples and not quantified at all in controls. The bars are coloured by cell type (FL83B and Hepa 1-6) and experimental condition (insulin stimulation versus basal). (**B**) Volcano plots of FL83B (upper row) and Hepa 1-6 (bottom row) samples before (left column) and after (right column) imputation. Y-axis denotes the log2 fold change after insulin stimulation, and the x-axis denotes the adjusted negative log10 p-value. The total number of differentially up- and down-regulated phosphosites are noted in each plot. Phosphosites were considered differentially regulated with the FDR adjusted p-value < 0.05.

**Figure S2.**
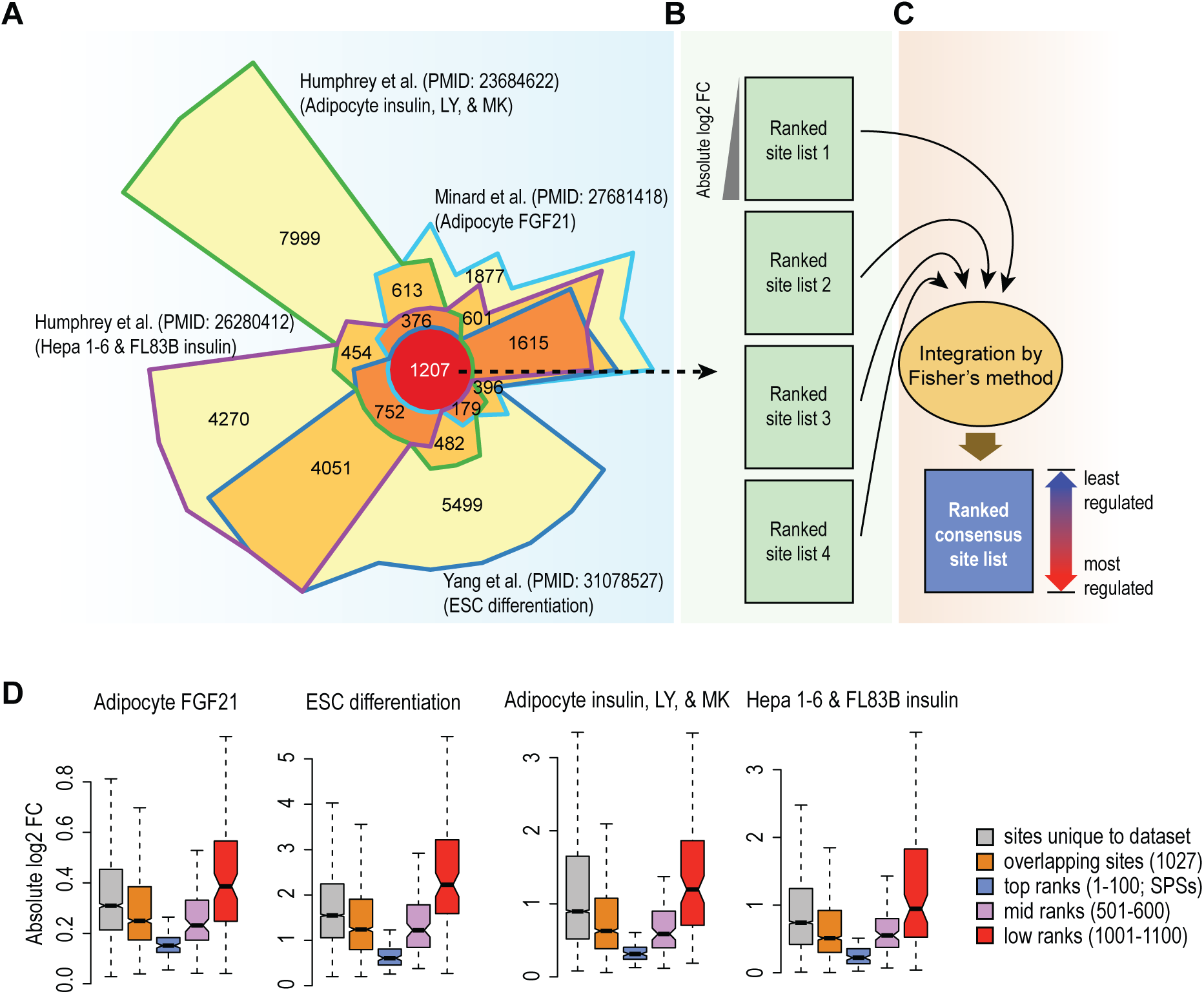
Identification of stably phosphorylated sites (SPSs) across phosphoproteomic datasets from different cell types and experimental conditions. (**A**) Chow-Ruskey diagram showing the four-way overlap of phosphorylation sites in four phosphoproteomic datasets generated from different cell types and under diverse experimental and cellular conditions. (**B**) A schematic depicting the ranking of phosphosites from low to high based on their absolute log2 fold change in each of the four datasets. (**C**) Integration of the individual rank lists using Fisher’s method for generating a consensus list of phosphosites in which they are ranked from least regulated to most regulated. (**D**) Boxplots of absolute log2 fold changes in phosphorylation between treatment and basal conditions. Phosphosites are grouped into five categories including those that are unique to each of the four phosphoproteomic datasets (gray), those that are found in at least two or three datasets (orange), and the top-ranked (blue), mid-ranked (purple), and low-ranked (red) phosphosites found in all four datasets. The common phosphosites were partitioned into the three groups on the basis of the consensus rank list generated in (**C**).

**Figure S3.**
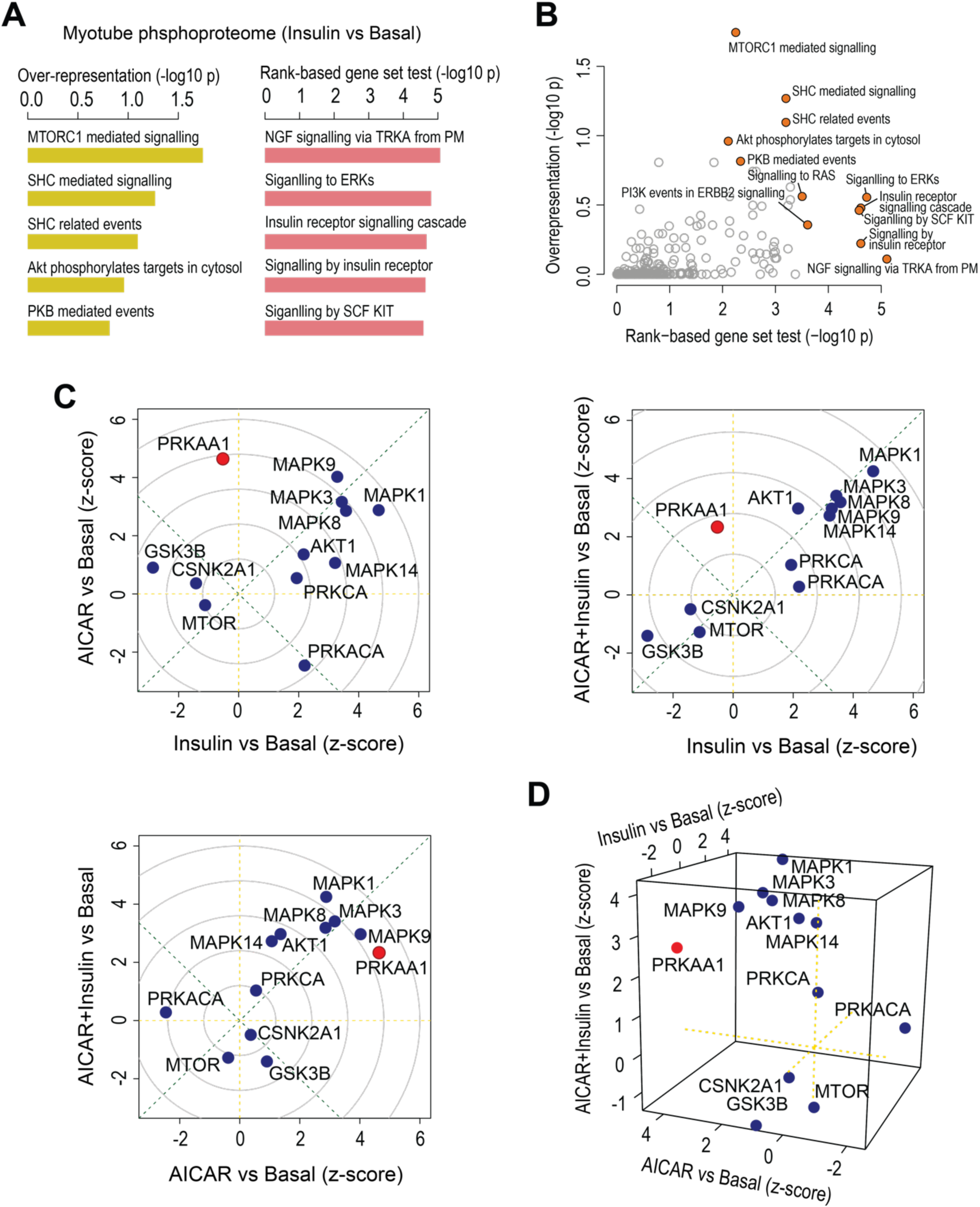
Demonstrations of 1-, 2-, and 3-dimensional gene-centric and site-centric analyses of the myotube phosphoproteome. (**A**) 1-dimensional (over-representation and rank-based gene set test) gene-centric pathway enrichment analysis of insulin versus basal conditions in myotube phosphoproteomic dataset. Reactome database was used for this enrichment analysis and the top most significant pathways are included in the barplot. (**B**) Scatter plot for comparing enrichment of pathways (in negative log10 p-values) between the two 1-dimensional pathway enrichment analysis. The x-axis and y-axis refer to the p-values derived from the rank-based gene set test and over-representation test, respectively. (**C**) 2-dimensional site-centric kinase activity analyses for all pairwise comparisons of the three treatments (AICAR, Insulin, AICAR+Insulin) versus basal. PRKKA1, a catalytic subunit of AMP-activated protein kinase (AMPK), is coloured in red. The x and y-axes denote the z-scores calculated for each kinase based on the overall changes of their substrates (phosphosites annotated to each kinase based on PhosphoSitePlus database) under each comparison. (**D**) 3-dimensional site-centric kinase activity analysis for comparing all three treatments in a single statistical analysis. The x, y, and z-axes denote the z-scores calculated of kinases for the three experimental conditions (Insulin vs Basal, AICAR vs Basal, and AICAR+Insulin vs Basal).

**Figure S4.**
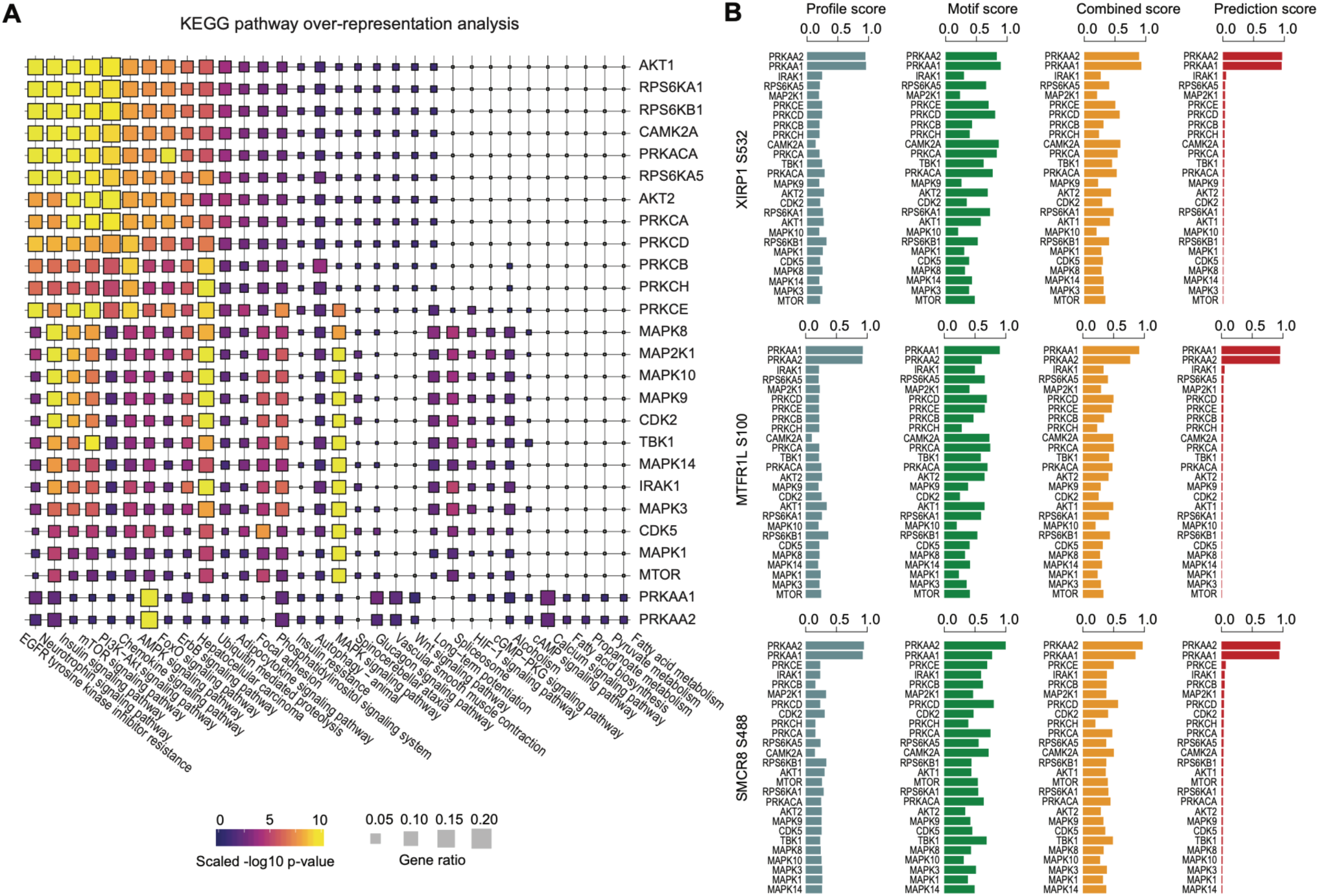
Pathway enrichment of kinase-substrates and kinase-substrate scores of example AMPK substrates. (**A**) KEGG pathway over-representation pathway analysis of kinase substrates (prediction score > 0.5). The squares are coloured by negative log10 *p*-value (scaled across pathways) and sized by the ratio of genes present in the KEGG gene set (the larger the square, the greater the proportion of kinase substrates present in the gene set). Any pathway that is not significantly represented is depicted as a grey square. Pathways were considered to be significantly over-represented when *p* < 0.05. (**B**) Bar plots of profile, motif, and combined scores, and positive-unlabelled ensemble learning prediction score of the top-ranked AMPK substrates.

**Figure S5.**
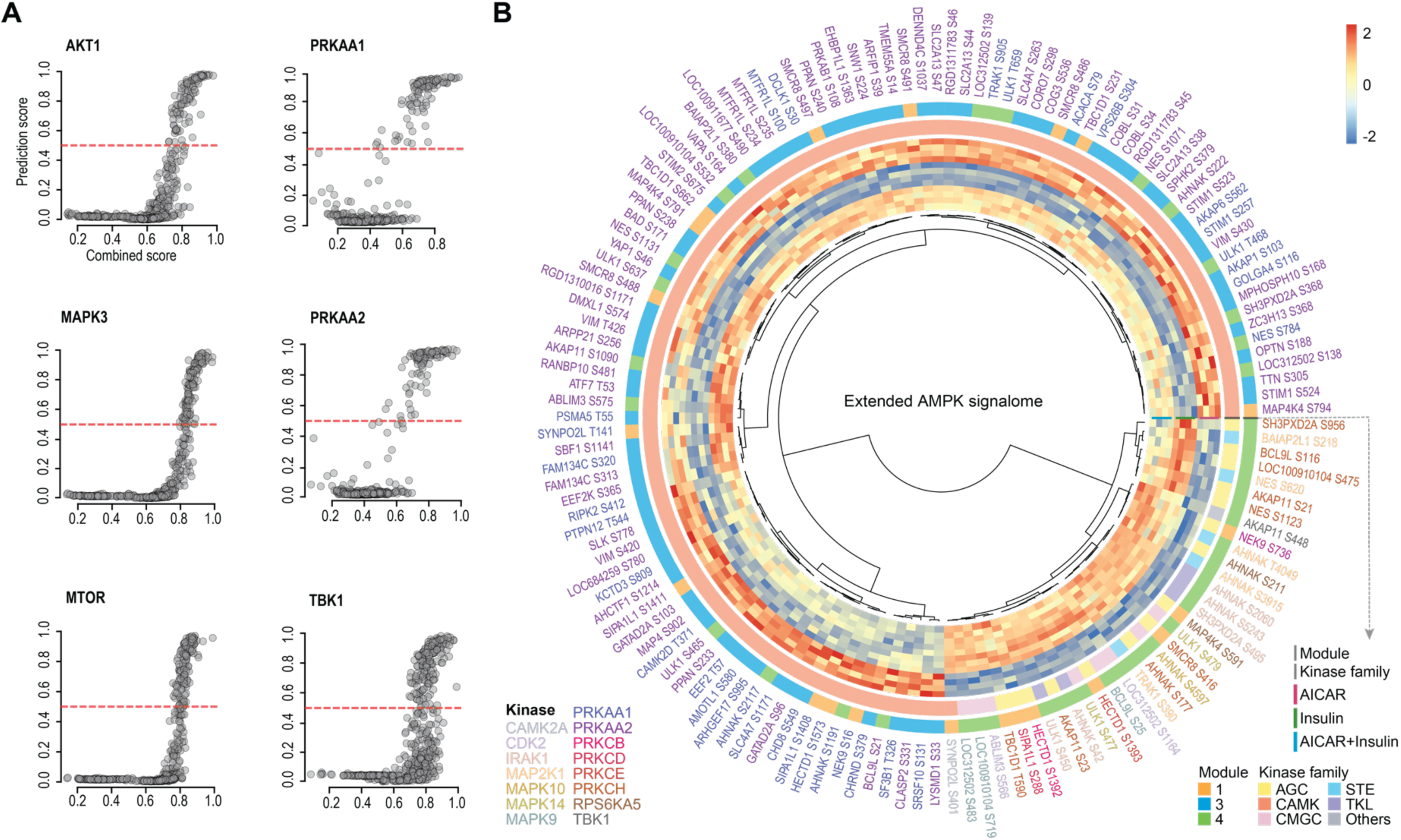
Generation and visualisation of signalomes. (**A**) Scatter plot of prediction score against combined scores of all identified phosphosites for AKT1, MAKP3, MTOR, PRKAA1, PRKAA2, and TBK1. (**B**) A circular heatmap representation of unsupervised clustering of the extended AMPK signalome. The phosphosites are coloured by their top kinase annotation, and the annotation bars highlighted by the protein module assignment of phosphosites (outer ring) and by kinase family of the kinases annotated for each phosphosite (inner ring).

**Figure S6.**
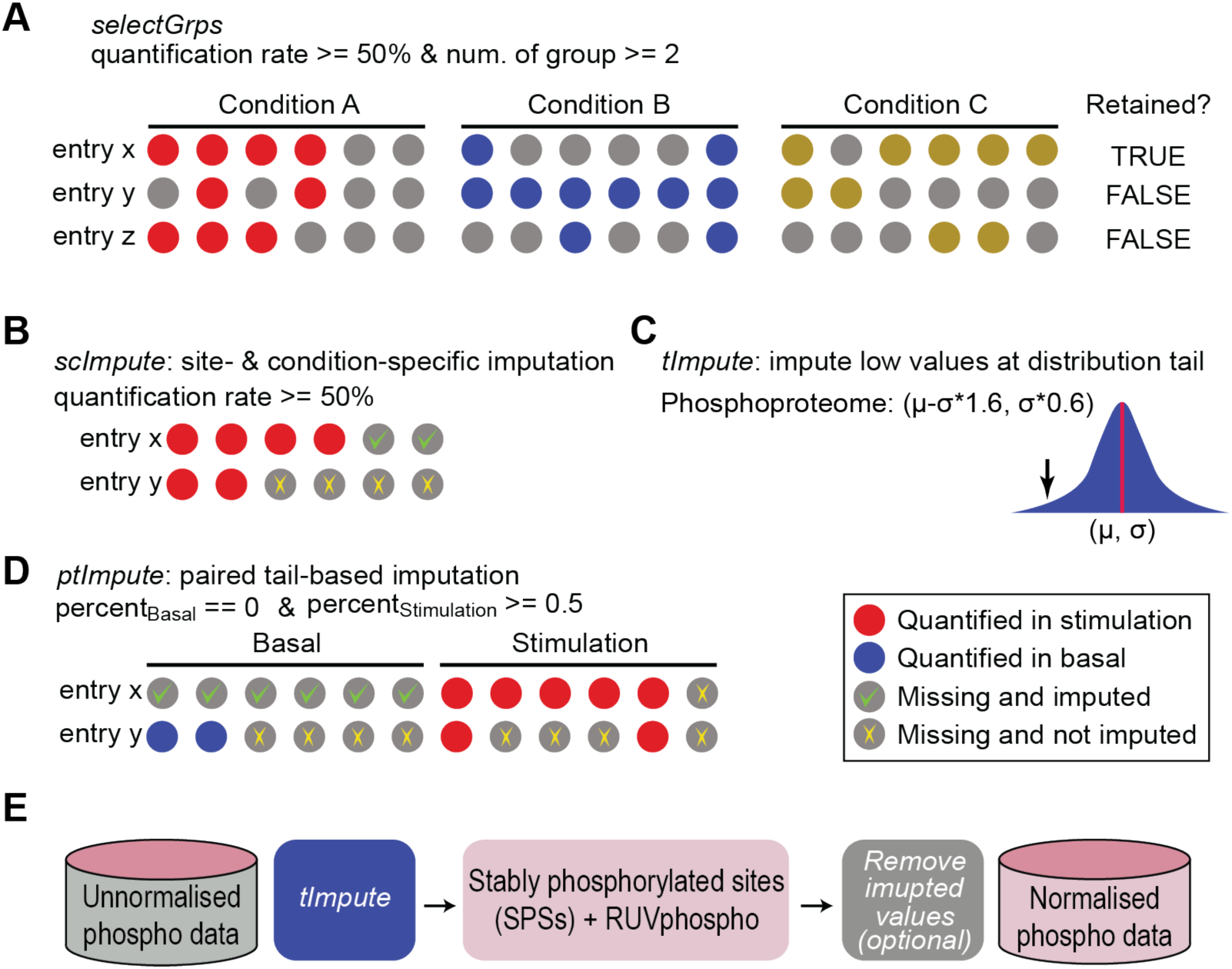
Schematic representation of the key filtering, imputation, and normalisation methods in PhosR. (**A**) An example of using *selectGrps* function to filter a phosphoproteomic data by selecting only rows (i.e., phosphosites) that have more than 50% of non-missing values (denoted as red, blue and gold circles) in at least two experimental groups (or conditions). (**B-D**) Illustration of the three imputation methods implemented in PhosR: *scImpute, tImpute*, and *ptImpute*. (**B**) An example of using *ssImpute* to impute missing values when more than 50% of replicates are quantified. (**C**) *tImpute* fits a distribution and imputes values from the distribution tail to mimic lowly expressed phosphosites. (**D**) *ptImpute* performs a paired tail-based imputation of regulated phosphosites. Here is an example that phosphosites detected in less than 50% of the “basal” or control samples and detected in greater than 50% of the “stimulated” or treated samples are selected for imputation, and vice versa for phosphosites detected in greater than 50% of the “basal” and in less than 50% of the “stimulated” samples. (**E**) A schematic representation of the batch correction framework implemented in PhosR with RUV normalisation using stably phosphorylated sites (SPSs) as negative controls. Missing data in phosphoproteomic data are imputed using *tImpute* and then normalised by using SPSs and removing unwanted variation (RUV). After normalisation, imputed data from *tImpute* are removed by default but can be retained by option of users.

